# Estimating the selection pressure and evolutionary rate of proteins on the non-neutral hypothesis of synonymous mutations

**DOI:** 10.1101/2024.04.14.589335

**Authors:** Jiachen Ye, Chunmei Cui, Rui Fan, Qinghua Cui

## Abstract

The Nonsynonymous/Synonymous substitution rate ratio (Ka/Ks) is a commonly used metric to estimate the selection pressure and evolutionary rate of proteins in comparative genomics, which plays critical roles in molecular evolution in both biology and medicine. A fundamental assumption of Ka/Ks is that synonymous mutations are evolutionarily neutral and not subject to natural selection as they do not alter protein sequences and function. However, a number of studies have demonstrated that synonymous mutations are non-neutral and may lead to diseases through a number of mechanisms, such as altering miRNA regulation. This further implies that synonymous mutations also participate in the process of natural selection and thus Ka/Ks should be redefined as well. For this purpose, here we propose an improved Ka/Ks ratio, *i*Ka/Ks, which re-computed the neutral substitution rate by taking the altered status of miRNA binding into consideration, and thereby incorporate the impact of synonymous mutations on miRNA regulation. As a result, *i*Ka/Ks shows better performance than Ka/Ks when comparing them using their correlation with expression distance. Moreover, case studies showed that *i*Ka/Ks is able to identify the positive/negative selection genes that are missed by Ka/Ks. For example, TMEM72/Tmem72 is estimated to be positively selected by *i*Ka/Ks (1.13) but negatively selected by the conventional Ka/Ks ratio (0.21). Further evidence showed its rapid evolution, which further support the power of the new algorithm.

## Introduction

The comprehension of species interrelations lies at the core of evolutionary biology^1^. Comparative genomics, as a cornerstone methodology in contemporary biomedical research^2^, involves scrutinizing genetic information within and across organisms to unveil the evolutionary trajectory, structure, and functionality of genes, proteins, and non-coding RNAs^3^. Through systematically comparative analysis, the biological relationships and evolutionary dynamics among species or among molecules can be methodically explored and evaluated. This facilitates a deeper understanding of the structure and function of orthologous genes, thus enriching our comprehension of human diseases and their potential therapeutic targets^4^.

Comparative genomics has provided a new means for understanding evolution by natural selection^5^. The ratio of nonsynonymous to synonymous substitution rate (Ka/Ks, also known as dN/dS) holds significantly importance for reconstructing phylogenies and elucidating the evolutionary dynamics of protein-coding sequences across closely related but diverged species^6,7^. The ratio serves as a pivotal metric for estimating the selection pressure and evolutionary rate of proteins and assessing the divergence among homologous gene sequences^8^. Generally, a diminished Ka/Ks value signifies heightened conservation within the homologous gene pair, with Ka/Ks << 1 indicating robust purifying selection. Conversely, an elevated Ka/Ks ratio signals increased disparity between homologous genes, with Ka/Ks > 1 suggestive of positive selection acting upon the gene. It is pertinent to note that the Ka/Ks algorithm is based on a fundamental assumption that synonymous mutations are neutral as they do not alter protein sequences and functions and thus are devoid of any influence on phenotypes and diseases^9^. According to this assumption, the synonymous substitution rate Ks functions as a baseline for the background nucleotide substitution rate, and the Ka/Ks ratio delineates the strength of natural selection driving gene evolution^10^.

However, increasing evidence has showed that synonymous mutations could be non-neutral^10,11^. They could even be considered to have comparable effects with nonsynonymous mutations in yeast^12^. Another study has brought to light the presence of codon usage bias at certain synonymous mutation sites, hinting at the potential non-neutrality of synonymous mutations and their consequential functional implications^13^. Notably, the association between miRNAs and synonymous mutations has emerged as a pivotal focus. This is because coding regions have been found to directly interact with miRNAs^14,15^ and a proportion (40%-60%) of miRNA binding sites have been identified to locate within mRNA coding regions^16,17^. Synonymous mutations may lead to melanoma^18^ and Crohn’s disease^19^ by altering miRNA binding. Wang *et al.* emphasized that nearly half of synonymous SNPs could modulate protein expression levels through miRNA-mediated gene regulation, leading to functional consequences and disease risks^20^.

These observations indicate that synonymous mutations have the potential to modify miRNA binding sites, consequently impacting gene function. Thus, assuming synonymous substitution rate as the background nucleotide substitution rate is questionable, and thus the Ka/Ks algorithm should be re-defined. For this purpose, here we present an improved Ka/Ks algorithm, *i*Ka/Ks, by considering the alteration of miRNA binding by synonymous mutations. The results showed that the new algorithm demonstrates better performance compared with the conventional one.

## Materials and methods

### Data collection

We obtained the coding sequences of all orthologous genes between human (GRCh38) and mouse (GRCm39) from Ensembl (https://www.ensembl.org/biomart/martview/)^21^, prioritizing the longest transcript sequence for genes with multiple transcripts. Subsequently, sequence alignment using MAFFT^22^ yielded a total of 22,106 pairs of orthologous genes. Mature miRNA sequences were retrieved from miRBase v22^23^.

### Evaluation of the miRNA-synonymy of mutation sites

Each mutant site is categorized as either synonymous or nonsynonymous based on whether it alters the corresponding amino acid sequence. Drawing a parallel to this classification, we distinguish between miRNA-synonymous and miRNA-nonsynonymous sites according to their influence on miRNA binding on target mRNAs. For each mutation, a sliding window of 31-nt is employed to capture the sequence of the site along with its flanking 15-nt on both sides. Subsequently, we predict the miRNA binding sites under four nucleotides (A,G,C,T) using TargetScan^24^. This prediction is facilitated by the Perl scripts of TargetScan 7.0 with default parameters. Therefore, for each mutation, we obtained two sets of miRNAs that regulate the mRNA respectively before and after the mutation. If these two sets are identical, the mutation is considered to be miRNA-synonymous; otherwise, it is considered to be miRNA-nonsynonymous.

### Calculation of *i*Ka/Ks of orthologous gene pairs

Following the methodology of KaKs Calculator 3.0^7^, for a pair of orthologous genes between human and mouse, insertion/deletion mutations were eliminated after sequence alignment using MAFFT. The number of nonsynonymous sites in the coding sequence (CDS) of the human gene is denoted as *N*, while the number of synonymous sites is denoted as *S*. Next, referencing the human gene, mutations in the mouse gene were tallied with *Nd* denoting the number of nonsynonymous mutations and *Sd* denoting the number of synonymous mutations. Consequently, the conventional Ka/Ks algorithm is as follows:

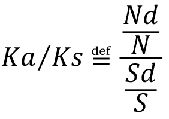

We next define the *i*Ka/Ks algorithm under the framework of miRNA regulation. Among *S* synonymous sites, the number of sites simultaneously considered as miRNA-synonymous is denoted as *Sm*. We consider these sites to be genuinely "neutral" implying they do not partake in any natural selection process. Similarly, among *Sd* synonymous mutations, the number of mutations simultaneously considered as miRNA-synonymous is donated as *Sdm*. Hence, the *i*Ka/Ks ratio is defined as follows:

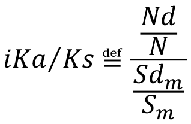

In summary, the essence of the improved algorithm lies in optimizing the background substitution rate to more accurately represent the random mutation rate unaffected by natural selection. The difference of these two algorithms is shown in Figure 1. The source code of the *i*Ka/Ks algorithm and the *i*Ka/Ks ratio data of the human-mouse orthologous genes are freely available at http://www.cuilab.cn/ikaks.

**Figure 1.**
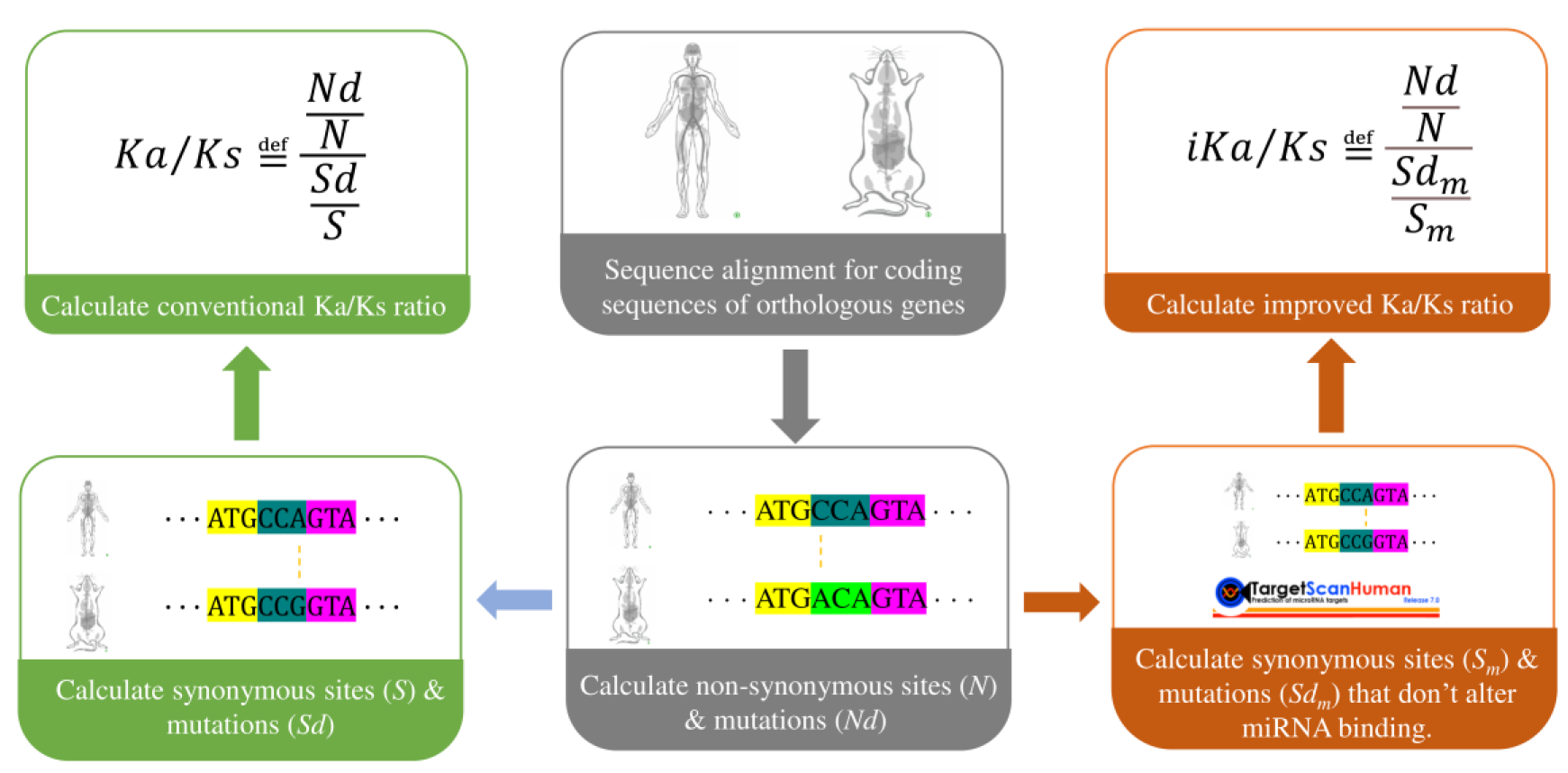
Workflow of the Ka/Ks algorithm and the *i*Ka/Ks algorithm.

### Comparison of *i*Ka/Ks and Ka/Ks ratios with gene expression distance

It is generally believed that the smaller the Ka/Ks ratio of homologous genes, the closer their expression profiles are. To authenticate the superiority of the improved algorithm, we obtained the expression profile data of orthologous genes for both human and mouse from Su *et al.’*s study^25^. This dataset comprises microarray expression profiles of 2275 orthologous gene-pairs across 27 corresponding tissues in human and mouse, including heart, kidney, lung, liver, cerebellum, testis and so on. Next, we used relative abundance (RA) to standardize the data, which can avoid errors stemming from variations in probe affinity for human and mouse tissues^26^. The RA for human or mouse gene *i* expressed in tissue *j* is defined as:

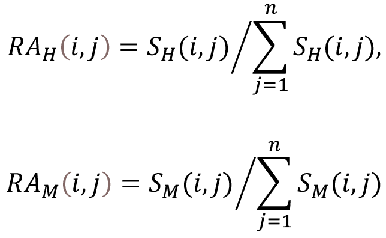

Here, *n* is the number of tissues considered as 27 in this study. *H* indicates human and M indicates mouse. 𝑆_𝐻_(𝑖, 𝑗) and 𝑆_𝑀_(𝑖, 𝑗) are the expression signal intensities of gene *i* in human tissue *j* and mouse tissue *j*, respectively. Subsequently, the Manhattan distance is used to define the expression distance of orthologous gene *i* between human and mouse:

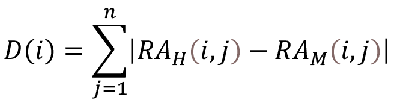

To validate the accuracy of the improved algorithm, we calculated the Spearman’s correlation between two Ka/Ks ratios and the expression distance *D(i)*, and then performed permutation tests by random assignment for 10000 times.

## Results

### The distribution of Ka/Ks and *i*Ka/Ks

For each human-mouse orthologous gene pair, we first counted the number of synonymous mutations affecting miRNA binding. The distribution of genes with synonymous mutations affecting miRNA binding is shown in Figure 2A. Among all the 22,106 orthologous gene pairs, half of them have over 100 synonymous mutations that can affect miRNA binding. This result suggests a large number of synonymous mutations potentially subject to natural selection through miRNA involvement, consistent with previous findings by Wang *et al* [15]. We next calculated the conventional Ka/Ks ratio and the *i*Ka/Ks ratio for each human-mouse orthologous gene pair. As a result, globally the *i*Ka/Ks ratio is highly correlated with the conventional one (R^2^=0.81, p-value=0.00). In the range where the Ka/Ks ratio is less than 0.4, their distributions are very close (D=0.0074, p-value=0.77, Kolmogorov-Smirnov test). However, as shown in Figure 2B, *i*Ka/Ks ratio is clearly distinguished from Ka/Ks for gene pairs with ratio>0.4 (D=0.16, p-value=1.17e-68, Kolmogorov-Smirnov test). It is noteworthy that among all the 22,106 orthologous gene pairs, *i*Ka/Ks exhibits 366 pairs with a ratio greater than 1, whereas the Ka/Ks only yields 11 pairs (OR=33.8, p-value=7.55e-94, Fisher’s Exact test). This indicates the potential presence of significant functional differences between orthologous genes of humans and mouse.

**Figure 2.**
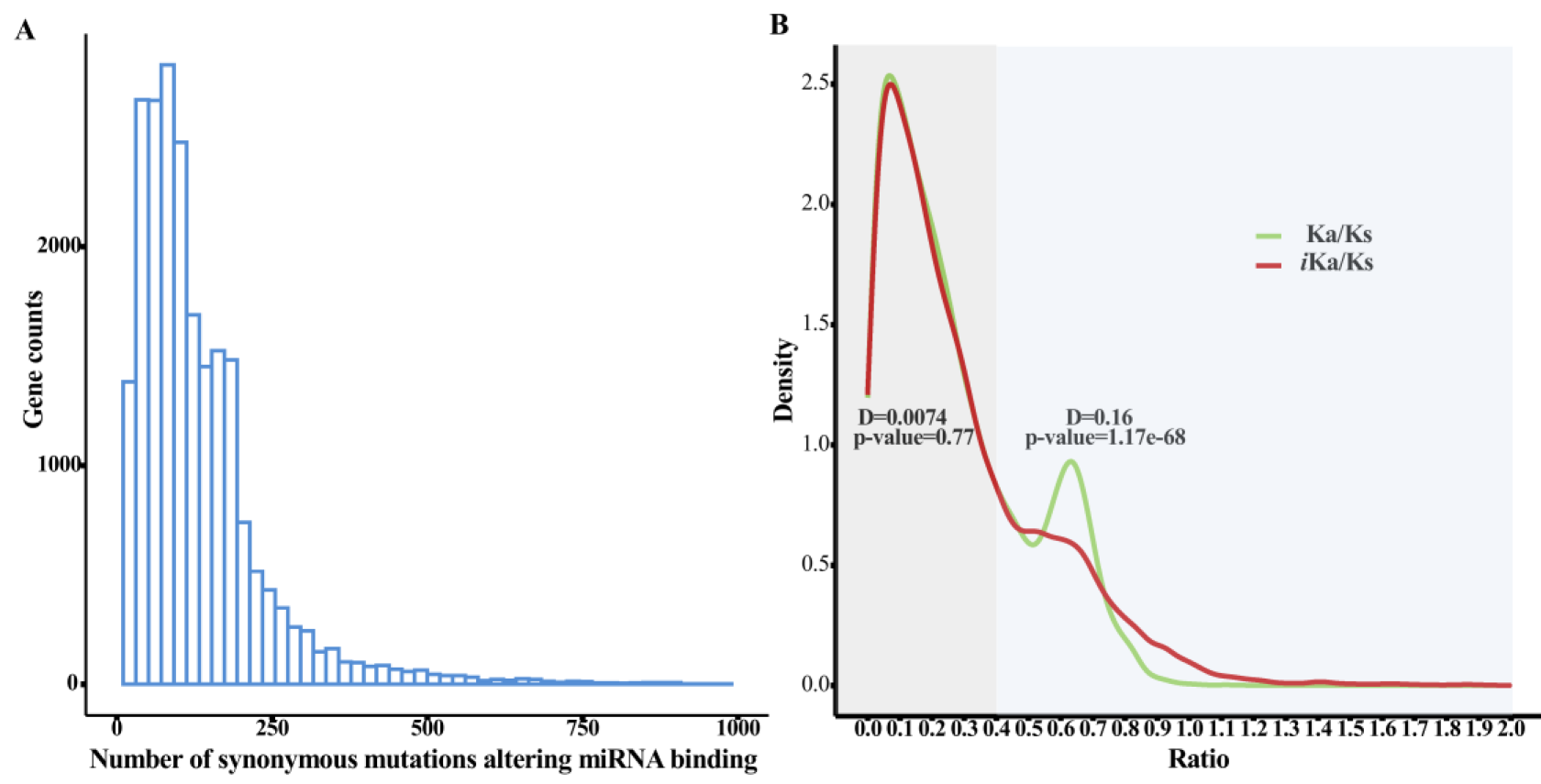
**(A) Statistics of the number of synonymous mutations altering miRNA binding.** **(B) Distributions of the Ka/Ks ratio and the *i*Ka/Ks ratio.** The Kolmogorov-Smirnov test is employed to assess differences between distributions. When the Ka/Ks ratio is less than 0.4, the distributions of the Ka/Ks ratio and the *i*Ka/Ks ratio are deemed highly similar (D=0.0074, p-value=0.77), whereas significant disparities between the distributions emerge when the ratio exceeds 0.4 (D=0.16, p-value=1.17e-68).

### The positive selection genes and the purifying selection genes identified by *i*Ka/Ks are consistent with the existing evidence

Among the 366 pairs of positive selection orthologous genes identified by iKa/Ks, 360 pairs have not been identified by conventional Ka/Ks. We first examined the cellular components of these positive selection genes in human using GO enrichment analysis. As a control, we also selected the top 500 pairs of orthologous genes with the largest differences between Ka/Ks ranking and *i*Ka/Ks ranking, simultaneously restricting *i*Ka/Ks to less than 0.1, as the purifying selection genes. The top 10 cellular components of these two groups of genes are shown in Figure 3A. The extracellular region and external side of plasma membrane are the primary locales for positive selection genes, while genes under purifying selection are predominantly situated within the nucleus, chromatin, and cytoplasm. Additionally, we observed keratin filaments as the top-ranked cellular component in the positive selection genes. This arises from the presence of numerous members of the KRTAP family (keratin associated proteins) among the positive selection genes identified by *i*Ka/Ks (Figure 3B). Importantly, several researches have reported rapid divergent evolution of KRTAPs between species^27,28^, which aligns with our findings. These findings illustrate that *i*Ka/Ks can identify positive selection genes as well as purifying selection genes missed by the conventional algorithms.

**Figure 3.**
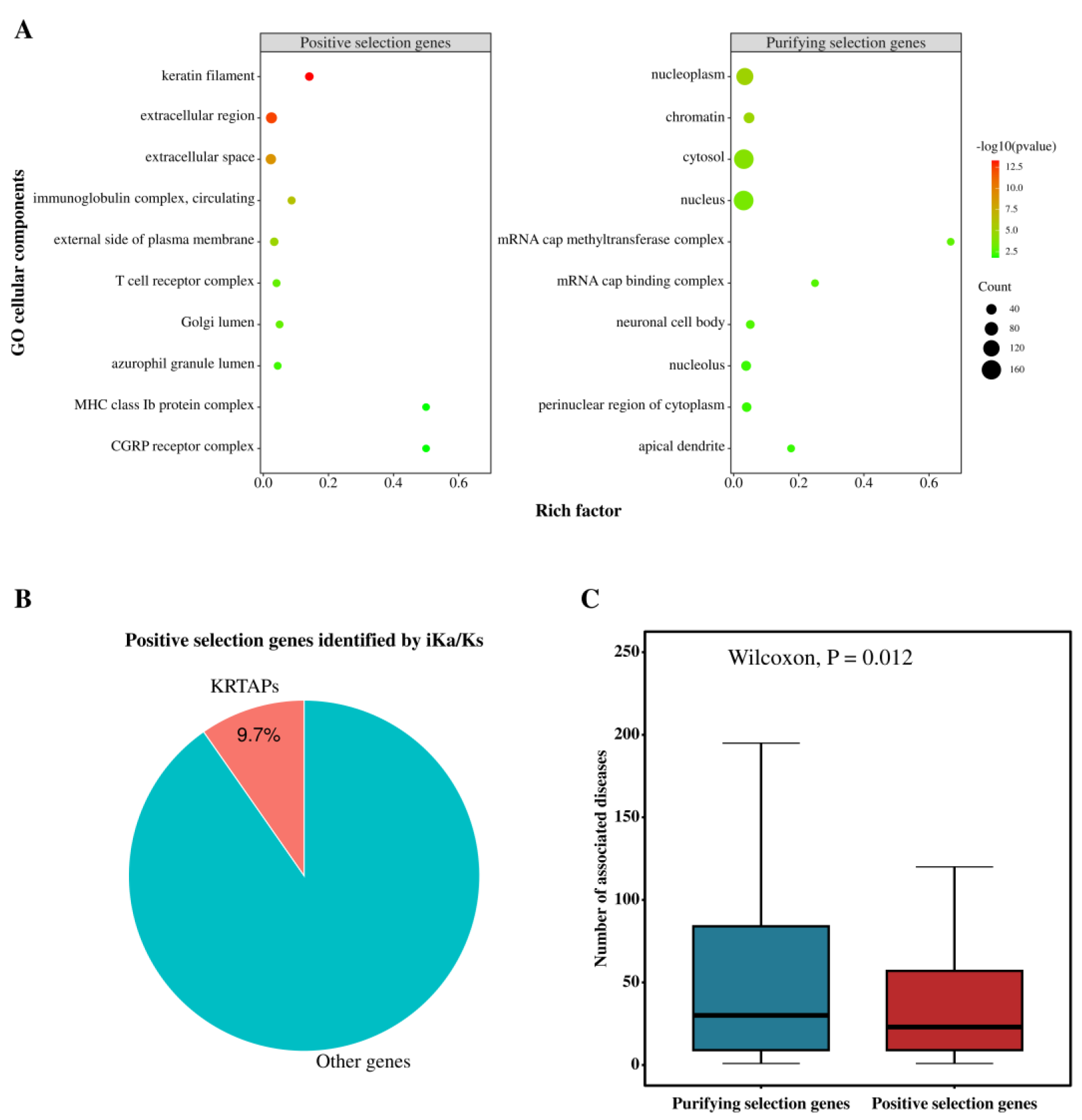
(A) Enrichment analysis of GO cellular components. The extracellular region and external side of plasma membrane are the primary locales for positive selection genes, while genes under purifying selection are predominantly situated within the nucleus, chromatin, and cytoplasm. Keratin filaments is the top-ranked cellular component in positive selection genes because of **(B) the presence of numerous members of KRTAPs**, which is considered as a rapid divergent evolution gene between species. (C) **Purifying selection genes exhibited significantly more associated diseases compared to positive selection genes (p-value=0.012, Wilcoxon Rank Sum test).**

Furthermore, we use DisGeNET^29^ to retrieve the number of human diseases associated with the genes and compared between the purifying selection genes and the positive selection genes. This result revealed that genes identified by *i*Ka/Ks undergoing purifying selection exhibited significantly more associated diseases compared to those undergoing positive selection (Figure 3C, Median value 30 vs. 23, p-value = 0.012, Wilcoxon Rank Sum test), further affirming the usefulness of *i*Ka/Ks.

### *i*Ka/Ks shows stronger correlation with expression distance compared to Ka/Ks

Next, we test whether the new algorithm outperforms the conventional one using their correlation with expression distance. It is widely hypothesized that homologous genes with notable sequence differences may exhibit considerable variations in expression. For doing so, we first obtained the gene expression profiles for 2275 pairs of homologous genes across 27 tissues in human and mouse. Subsequently, we calculated their expression distance, herein defined as the relative abundance variances across distinct tissues.

We first investigated whether the new algorithm outperforms the conventional algorithm within gene sets characterized by the maximum and the minimum expression distance. As shown in Figure 4A, it is obvious that genes with smaller expression distance have significantly smaller value of *i*Ka/Ks ratio compared to Ka/Ks ratio (Median value 0.097 *vs.* 0.093, p-value=0.029, Wilcoxon Signed Rank test). While the top 20% genes with larger expression distance exhibits a significantly greater value of *i*Ka/Ks ratio compared to Ka/Ks (Median value 0.20 *vs.* 0.21, p-value=0.012, Wilcoxon Signed Rank test). These results indicates that *i*Ka/Ks can better reflect the expression distance of orthologous genes than the conventional one.

**Figure 4.**
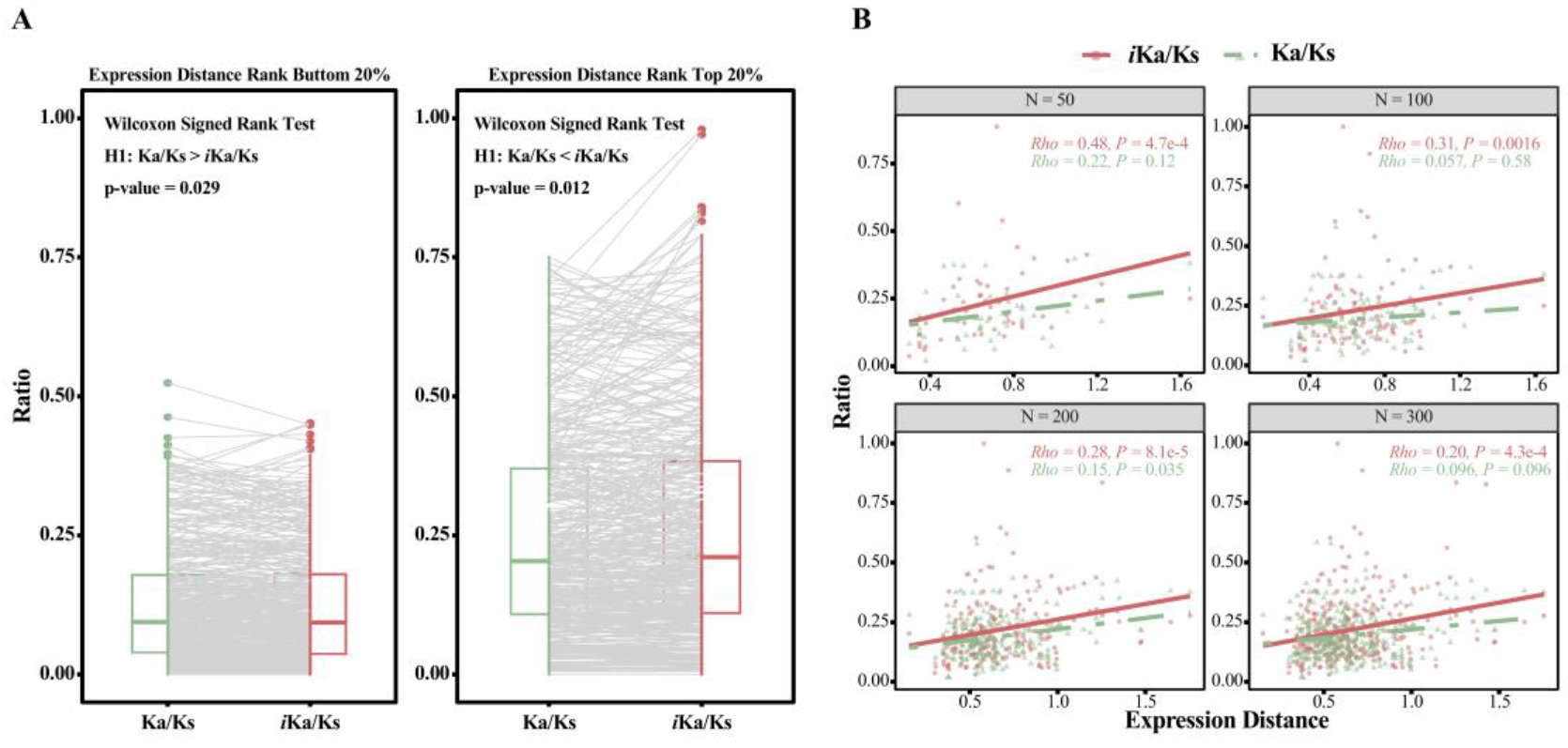
(A) Boxplot comparing the two algorithms grouped by the lowest and highest expression distance. In the group representing smaller expression distance, *i*Ka/Ks yields a smaller ratio (Median value 0.097 *vs.* 0.093, p-value=0.029, Wilcoxon Signed Rank test). Similarly, in the group representing larger expression distance, *i*Ka/Ks yields a larger ratio (Median value 0.20 *vs.* 0.21, p-value=0.012, Wilcoxon Signed Rank test). **(B) Correlation between *i*Ka/Ks or Ka/Ks and the expression distance.** For top *N* gene pairs with the greatest rank differences between the two algorithms, *i*Ka/Ks shows higher Rho and lower p- value (Rho=0.48, 0.31, 0.28, 0.20 *vs.* 0.22, 0.057, 0.15, 0.096; p-value=4.7e-4, 1.6e-3, 8.1e-5, 4.3e-4 *vs.* 0.12, 0.58, 0.035, 0.096 for *N*=50, 100, 200, 300, respectively).

Furthermore, we investigated whether the new algorithm exhibits a stronger correlation with expression distance compared to the conventional algorithm. Given that *i*Ka/Ks ratio is globally highly correlated with Ka/Ks, here we only focused on genes with highest difference between these two algorithms. Thus, we ranked genes by their Ka/Ks values within each algorithm and selected the top *N* genes with the greatest rank differences (here *N*=50, 100, 200, 300, respectively). Subsequently, correlation analysis between the *i*Ka/Ks or Ka/Ks ratio and the expression distance was performed within these four groups. As a result, *i*Ka/Ks showed higher correlation with the expression distance than Ka/Ks across all of the four groups (Figure 4B, Rho=0.48, 0.31, 0.28, 0.20 *vs.* 0.22, 0.057, 0.15, 0.096, p-value=4.7e-4, 1.6e-3, 8.1e-5, 4.3e-4 *vs.* 0.12, 0.58, 0.035, 0.096 for *N*=50, 100, 200, 300, respectively, Spearman’s correlation). For the groups for *N*=50, 100, and 300, the conventional algorithm didn’t even achieve statistical significance (with p-values of 0.12, 0.58, and 0.096, respectively). This result indicates that the new algorithm exhibits a significantly stronger correlation with expression distance than the conventional algorithm. To comprehensively test the significance of *i*Ka/Ks algorithm, we further performed permutation tests by random assignment. The result indicated that *i*Ka/Ks shows significantly higher correlation with expression distance than the conventional ratio (Supplementary Figure S1, p-value=0.0798, 0.0052, 0.0175, 0.0081 for *N*=50, 100, 200, 300, respectively. Permutation test for 10000 times). These results together suggest that the new algorithm outperforms the conventional one in this expression distance comparison.

## Case studies

By several case studies, here we showed that *i*Ka/Ks algorithm is able to identify positively selected genes that were estimated to be under purifying selection by conventional Ka/Ks ratio. For example for the human-mouse orthologous transmembrane protein 72 gene *TMEM72* and *Tmem72* ^30^, its conventional Ka/Ks ratio is 0.21, whereas its *i*Ka/Ks is 1.13, ranking 21,903 out of 22,106 gene pairs (Supplementary Figure S2A-B). This indicates a relatively poor conservation of this homologous gene between human and mouse, and suggests positive selection during evolution and potentially substantial functional divergence. According to GENEVESTIGATOR^®31^, the human *TMEM72* is specifically expressed in the kidney and rectum, with no significant expression in the nervous system as shown in Supplementary Figure S3. In contrast, as shown in Figure S4, its mouse orthologous is highly expressed not only in kidney but also in the nervous system, including the lumbar spinal ganglion, brain microvessel and the cerebellum *etc*. To further validate this finding, we employed the cross-species functional divergence analysis proposed by Cui *et al*. ^32^, which defined the functional divergence score (FDS) for orthologous genes between human and mouse and is able to discover conserved functions and species-specific functions of the orthologous genes. As a result, this gene has functions of cell migration, osteoblast differentiation, and vascular remodeling in both human and mouse (Figure 5). However, it has much more specific functions in human or in mouse. It shows functions on nervous system including neuron development and neuron apoptosis only in mouse but not in human, which is consistent with our findings above. Furthermore, it shows functions of latent virus replication and antiviral immunity *etc*. only in human but not in mouse. This significant divergence in expression pattern and in functions suggest that this gene could be indeed under positive selection.

**Figure 5.**
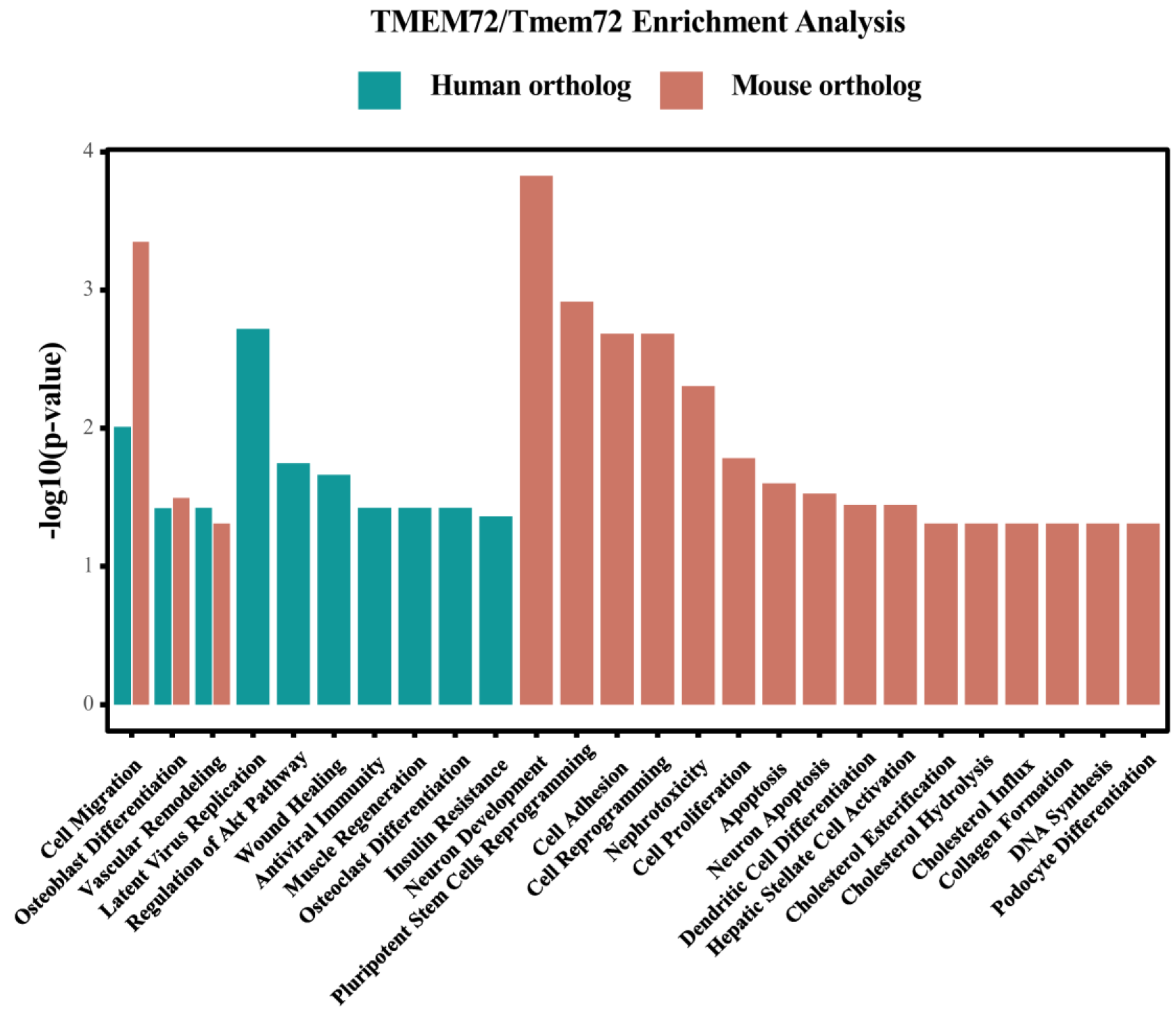
Functional Enrichment Analysis of TMEM72/Tmem72. This gene has functions of cell migration, osteoblast differentiation, and vascular remodeling in both human and mouse. Functions on nervous system including neuron development and neuron apoptosis are only found in mouse but not in human. And functions of latent virus replication and antiviral immunity *etc*. are only found in human but not in mouse.

Similarly, there exist another pair of orthologous genes, *RAMP1/Ramp1*, whose *i*Ka/Ks ratio (1.05) is also much greater than Ka/Ks ratio (0.42). It was reported by NCBI that RAMP1 is a member of the single transmembrane domain protein family known as the RAMP, possessing an extracellular N-terminus and a cytoplasmic C-terminus. According to the DEGs^33^, this gene is essential in human pluripotent stem cells, whereas gene knockout experiments in mice from the MGI^34^ database and phenotype analysis from IMPC(www.mousephenotype.org)^35^ suggest that its orthologous in mouse is non-essential. This differential essentiality further means functional divergence of this gene between human and mouse. Indeed, the functional divergence analysis revealed that the FDS score of *RAMP1/Ramp1* is 0.89, ranking 13,979 out of total 15,505 pairs, indicating that this pair of orthologous genes is highly diverged in function between human and mouse. This result suggests that this gene may be indeed under positive selection, which also supports the *i*Ka/Ks algorithm.

The new algorithm not only can identify the homologous gene pairs with significant differences (as shown in the above two case studies) but also is able to reveal functionally conserved orthologous genes between human and mouse. For instance, the *i*Ka/Ks ratio of the *IL1B/Il1b* orthologs is 0.23, which is smaller than conventional Ka/Ks ratio (0.42). The protein encoded by this gene belongs to the interleukin-1 (IL-1) cytokine family. The IL-1 cytokine is an important mediator of the inflammatory response, and is involved in a variety of cellular activities, including cell proliferation, differentiation, and apoptosis^36,37^. According to DEGs and MGI, its orthologues in human and mouse are both essential genes. Furthermore, according to DisGeNET, it is associated with 1801 diseases. Clearly, this gene plays a crucial role in the growth and development processes of both human and mouse, exhibiting a high degree of conservation, which is consistent with the analysis results and further supports *i*Ka/Ks algorithm.

## Discussion

Comparative genomics has yielded substantial insights in biology and medicine. Among several animal models, mouse stands out as the preferred model organism for studying human diseases owing to their compact size, docile disposition, swift reproductive cycle, and cost- effectiveness in maintenance^38^. Therefore, the similarity of orthologous genes between human and mouse serves as critical information, influencing the translatability of experiments conducted in mouse models^39^. While the Ka/Ks ratio has traditionally been a widely referenced metric for measuring evolutionary rate and selective pressure in comparative or evolutionary genomics One basic assumption of the Ka/Ks algorithm is that synonymous mutations are evolutionarily neutral, which, however, has recently been found to be inaccurate, that is a large number of synonymous mutations could be not neutral but functional Based on the above observation, this study proposes a new Ka/Ks algorithm, *i*Ka/Ks, specifically focusing on the impact of synonymous mutations on miRNA binding sites. The results showed that the algorithm can more accurately assess the functional divergence, evolutionary rate and selective pressure of the orthologous genes between human and mouse. Specifically, the new algorithm demonstrates a stronger correlation with the divergence of gene expression patterns between human and mouse.

Furthermore, evidence supports that *i*Ka/Ks rectifies previous misidentifications of genes as either conservative or non-conservative. For instance, the *TMEM72/Tmem72* pair, is previously considered to be under purifying selection, shows an *i*Ka/Ks ratio > 1, indicating substantial divergence consistent with additional evidence.

In the future, this study could be continuously improved. While the *i*Ka/Ks ratio primarily focuses on sequence-level disparities in the coding region of orthologous genes, it is essential to recognize that other factors contribute to functional divergence among these genes. Research suggests that epigenetic modifications^40^, mutations in non-coding regulatory elements^41^, and the regulation of three-dimensional genome structure^42,43^ also play significant roles. Future studies may aim for comprehensive analyses that integrate information from upstream and downstream regions of the entire genome, as well as structural insights, to gain a deeper understanding of the functional disparities among orthologous genes. In addition, applying *i*Ka/Ks to SNPs among human populations may help to find some important mutation sites playing important roles in human phenotype and disease.

## Acknowledgements

This study was supported by the grants from the National Natural Science Foundation of China [62025102, 32301239, 81921001] and the Scientific and Technological Research Project of Xinjiang Production and Construction Corps [2023AB057].

## Author Contributions

Q.C. proposed the original idea and supervised the study. J.Y. performed the study. C.C. and R.F. provided helps in implementing the new algorithm. J.Y. wrote the raw manuscript and Q.C. edited the manuscript.

---

**Figure S1.**
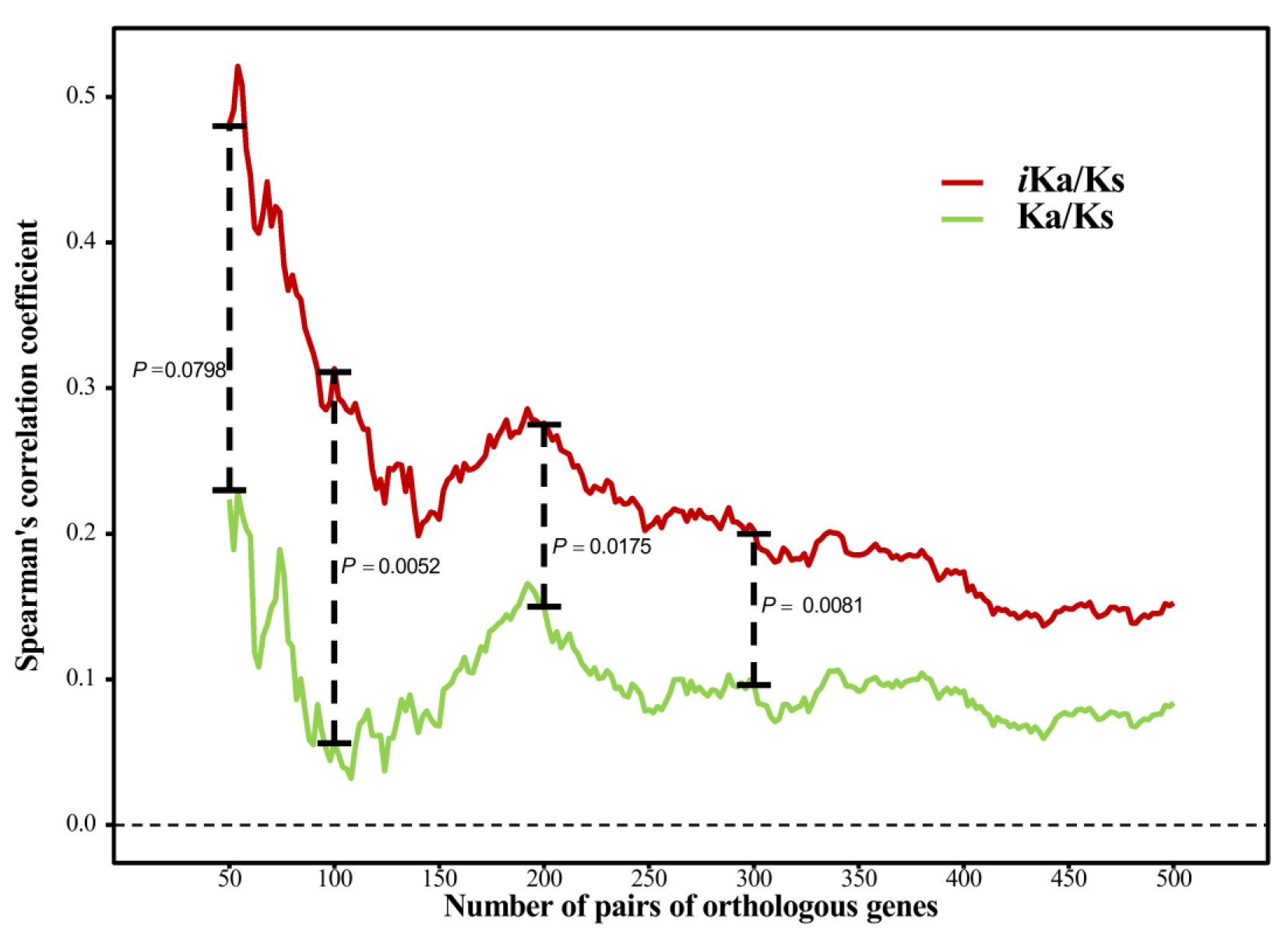
Correlation between *i*Ka/Ks or Ka/Ks and expression distance for *N*∈ [50,500]. The spearman’s correlation coefficient with expression distance of *i*Ka/Ks is higher than that of Ka/Ks across *N*∈[50,500]. After randomizing and partitioning two sets of data, this process was repeated 10,000 times to test how often a greater difference in correlation coefficients could be obtained compared to the original arrangement (798, 52, 175, 81 times for *N*=50,100,200,300, respectively). This permutation test aimed to assess the significance of the superiority of the *i*Ka/Ks algorithm.

**Figure S2.**
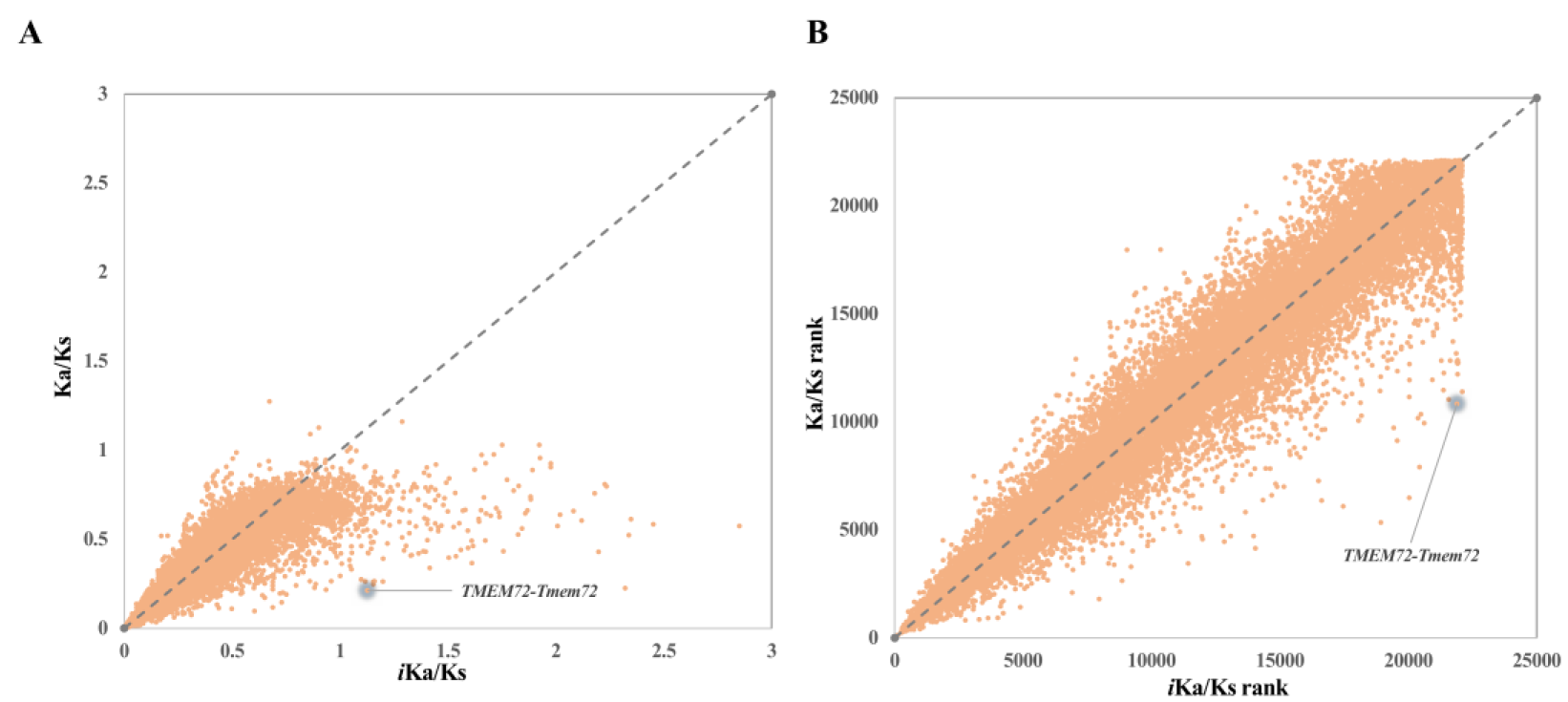
(A) The Ka/Ks ratio and the *i*Ka/Ks ratio of all gene pairs and the gene pair of *TMEM72-Tmem72*. (B) Rank of the Ka/Ks ratio and the *i*Ka/Ks ratio of all gene pairs and the gene pair of *TMEM72-Tmem72*.

**Figure S3.**
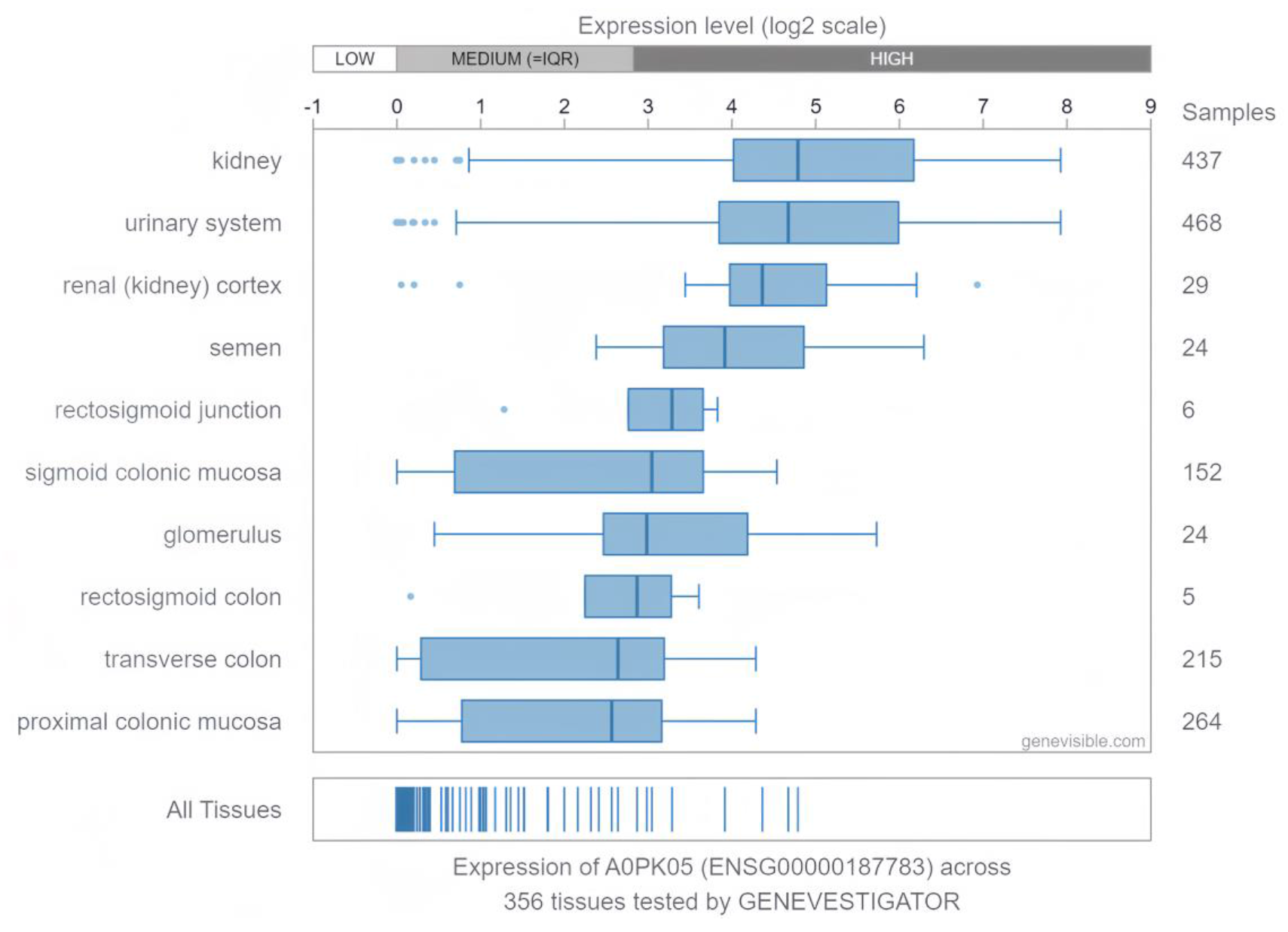
The expression profile of TMEM72 across 356 human tissues.

**Figure S4.**
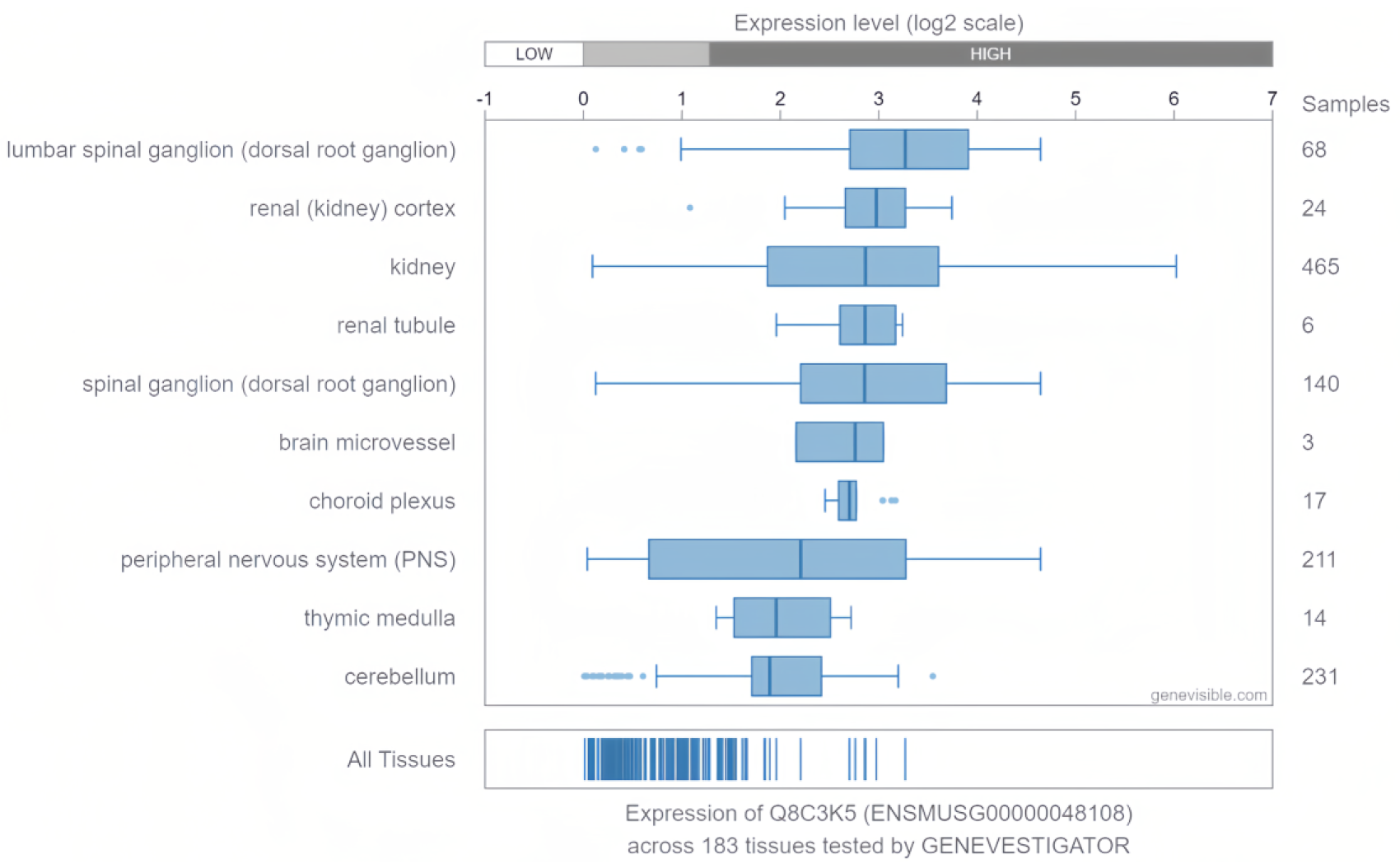
The expression profile of Tmem72 across 183 mouse tissues.

